# Pathogen-associated odor experience induces inherited olfactory avoidance in a parasitoid wasp

**DOI:** 10.64898/2026.09.11.751080

**Authors:** Liu Shiyao, Ge Jin, Sun Jianghua

## Abstract

Odors allow animals to evaluate resources and risks, but the neural and inherited consequences of experience with ecologically meaningful odors remain poorly understood. Since both parasitoids and entomopathogenic fungi are widely used as agents in integrated biocontrol, we asked whether exposure to pathogen-associated volatiles can reassign odor value in a parasitoid wasp and whether the acquired bias is transmitted to descendants. Adult female *Scleroderma guani* were exposed, without host contact, to volatiles from *Beauveria bassiana*-infected hosts or to methyl cinnamate (MC), a representative component of the infected-host odor. Twelve-hour exposure shifted subsequent choices away from infected-host odor and MC, whereas shorter exposure produced different effects, indicating duration-dependent olfactory plasticity. The shift occurred without an explicit reward or punishment and is therefore consistent with non-associative learning. Unexposed F1 and F2 descendants of MC-exposed females expressed the same avoidance bias. MC stimulation elicited stronger calcium responses in mushroom-body regions of trained females and their descendants. Head transcriptomes identified reduced expression of the crustacean cardioactive peptide receptor (CCAPR) as a shared molecular signature, and CCAPR RNA interference phenocopied the odor-avoidance behavior. Finally, maternal treatment with sinefungin, a broad S-adenosylmethionine-dependent methyltransferase inhibitor, left maternal behavioral shift intact but abolished the F1 bias, implicating a methylation-sensitive process in transmission. Together, these results reveal that a pathogen-associated odor can induce durable, cross-generational reweighting of olfactory value in a non-model insect and identify candidate neural and neuropeptidergic mechanisms linking sensory experience to inherited behavior, and providing a new perspective on utilizing training in integrated biocontrol as well.

## Introduction

Odors do more than identify objects: they carry values that determine whether an animal approaches, ignores, or avoids a stimulus. Those values are partly innate but can be recalibrated when experience changes the predicted consequences of a cue. Such flexibility is especially important when the same resource can shift between profitable and dangerous states. In insects, odor information is transformed across a hierarchy extending from olfactory receptor neurons and the antennal lobe to higher-order centers, including the mushroom body and lateral horn, where sensory identity is integrated with experience and behavioral state (Masse et al., 2009; Perry et al., 2013; Schultzhaus et al., 2017).

Experience-dependent changes in odor-guided behavior do not always require explicit pairing between an odor and a reward or punishment. Repeated or prolonged stimulation can produce non-associative plasticity, classically described as habituation or sensitization, and can alter behavior at peripheral, central, or both levels of the olfactory pathway (Perry et al., 2013). In insects, acute sensory exposure can generate long-lasting changes in odor responses, whereas chronic exposure at ecologically relevant concentrations may produce limited peripheral plasticity and stronger changes downstream (Anton et al., 2011; Gugel et al., 2023; Minoli et al., 2012). Parasitoid wasps provide rich examples of experience-dependent host searching, but most work has emphasized associative learning during host contact, feeding, or oviposition (Giunti et al., 2015; Haverkamp and Smid, 2020).

A further unresolved question is whether the consequences of sensory experience end with the individual. Parental olfactory experience can bias odor-guided behavior in descendants of flies, butterflies, nematodes, and mammals (Bozler et al., 2019; Dias and Ressler, 2014; Gowri et al., 2019; Remy, 2010; Williams, 2016). Recent experiments in butterflies further suggest that circulating factors, but not the odorant itself, can contribute to the transmission of acquired preferences (Chua et al., 2026). Nevertheless, mechanistic links among parental sensory experience, altered neural processing in descendants, and germline or maternal transmission remain rare, particularly in ecologically specialized non-model insects.

Parasitoids offer a stringent ecological test of this problem because the value of a host depends on its physiological state. Very often parasitoids and microbial agents are jointly applied in Integrated Pest Management (IPM), however, infection by entomopathogenic fungi alters host-associated odors and can sharply reduce host suitability for parasitoid development (Roy and Pell, 2000; Koller et al., 2023; Pobozniak and Olczyk, 2025). Fungal associates and fungal infection can therefore change host location and acceptance by parasitoids (Mesquita and Lacey, 2001; Sullivan and Berisford, 2004; Miranda-Fuentes et al., 2021; Wu et al., 2022). These interactions are often discussed as a problem of biocontrol compatibility (Quesada-Moraga et al., 2022; Koller et al., 2023), but they also raise a broader biological question: can experience with a pathogen-associated odor durably change how an animal assigns value to that cue, and can this change be inherited?

We addressed this question in the bethylid wasp *Scleroderma guani*, an ectoparasitoid of wood-boring beetles. *S. guani* serves as a main parasitoid against *Monochamus alternatus,* the vector of pinewood nematode that causes the pine wilt disease (Xu et al., 2008; Zhao et al., 2014). The entomopathogenic fungus *Beauveria bassiana,* a common biocontrol agent widely distributed in nature, resulted the infected *Monochamus alternatus* hosts a poor reproductive resource for the wasp (Chen et al., 2026; Zimmermann, 2007). Meanwhile, methyl cinnamate (MC) is a prominent component of the infected-host volatile blend recognized by *S. guani* (Huang et al., 2025). We asked whether odor exposure alone changes subsequent odor choice, whether the resulting bias persists in unexposed F1 and F2 descendants, and whether the phenotype is associated with changes in mushroom-body activity, neuropeptide signaling, and methylation-dependent transmission. By separating these questions from the applied context, the study tests a general mechanism by which ecologically meaningful sensory experience can reshape behavior within and across generations.

## Results

### Prolonged exposure to pathogen-associated odor reverses odor preference without host contact

Naive females showed little discrimination between odors from healthy and *B. bassiana*-infected hosts (Fig. 1 A). After 12 hours of direct exposure to an infected host, females subsequently preferred healthy-host odor. The same shift occurred when females were physically separated from the infected host and experienced only its volatiles, demonstrating that host contact, feeding, and oviposition were not required (Fig. 1 A and B). Exposure to healthy-host odor also increased subsequent preference for healthy hosts, indicating that experience altered the weighting of host-associated cues rather than requiring direct experience of fungal pathology.

**Figure 1.**
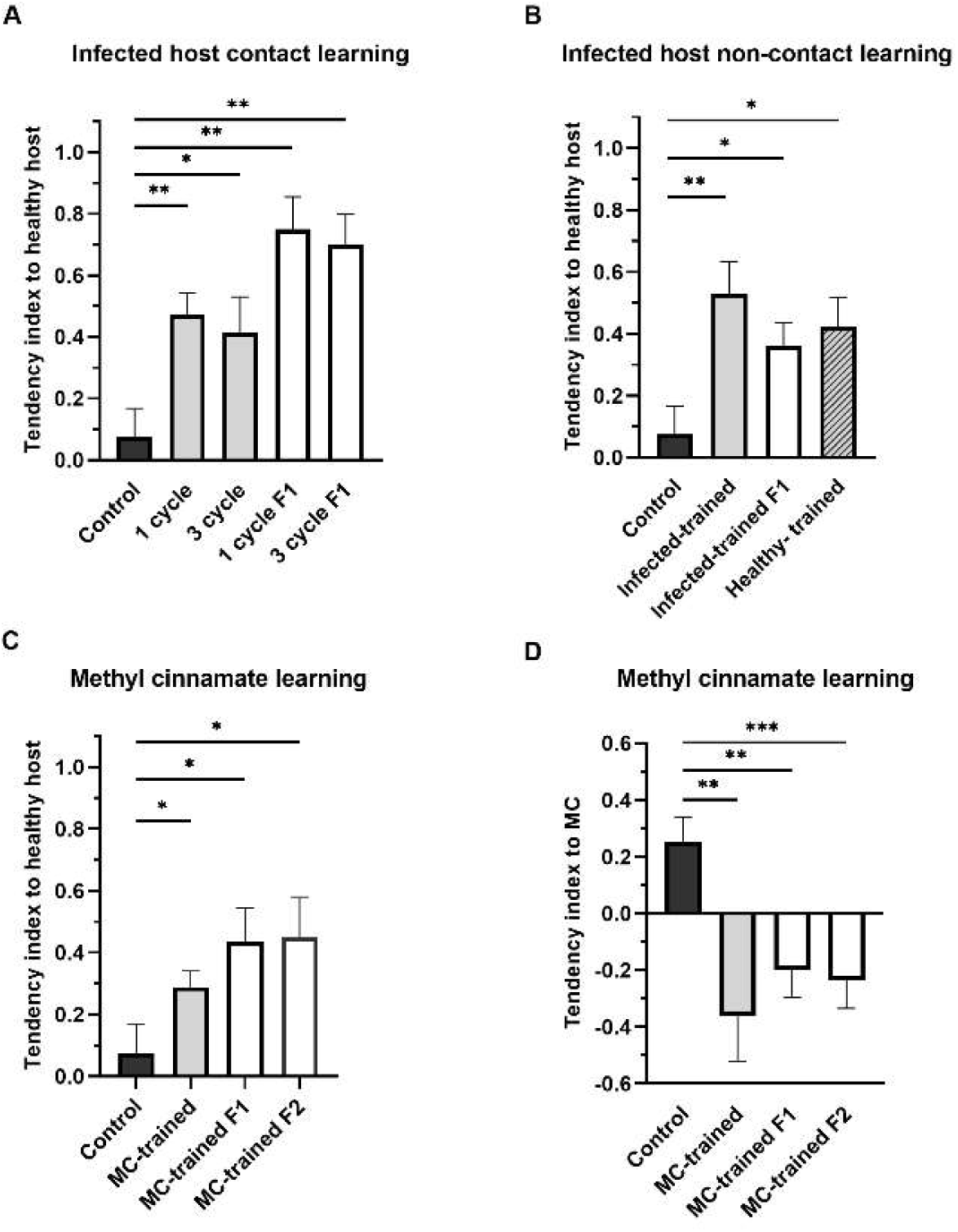
Experience-dependent and inherited changes in odor preference. (A) Preference after direct contact with infected hosts and in F1 descendants. (B) Preference after noncontact exposure to infected-host volatiles and in F1 descendants. (C) Preference for healthy versus infected hosts after MC exposure and in F1 and F2 descendants. (D) Preference for MC versus solvent after MC exposure and in F1 and F2 descendants. MC, methyl cinnamate. Data are means ± SE. ns, not significant; *P < 0.05; P < 0.01;* P < 0.001.

We next isolated MC as a representative component of the infected-host blend. Twelve-hour exposure to MC reduced subsequent choice of both MC and infected-host odor (Fig. 1 C and D). The effect depended on exposure duration: a short 0.5-h exposure increased attraction to infected-host odor, whereas longer exposures progressively shifted behavior toward avoidance (Fig. S4). Thus, odor experience did not produce a simple monotonic loss of responsiveness but a duration-dependent reassignment of cue value. Because the odor was presented without an explicit reinforcing event, the behavioral change is consistent with non-associative olfactory plasticity.

### Maternal odor experience biases odor choice in F1 and F2 descendants

To test whether the acquired bias extended beyond the exposed generation, trained females were allowed to reproduce without further behavioral testing. Unexposed F1 offspring of females trained by infected-host contact or by infected-host volatiles showed the same preference for healthy-host odor as their mothers (Fig. 1 A and B). Repeating the contact experience did not further strengthen the F1 phenotype.

The effect of MC exposure persisted for at least two generations. Unexposed F1 and F2 descendants of MC-trained females showed reduced preference for MC and increased preference for healthy-host odor relative to descendants of naive females (Fig. 1 C and D). Under this maternal exposure design, the F1 effect is intergenerational, whereas persistence in F2 supports a transgenerational component.

### Trained females and their descendants show enhanced mushroom-body responses to MC

We used histological sections to identify the antennal lobes, central brain, and mushroom bodies of *S. guani*, and then recorded odor-evoked calcium signals from semi-intact head preparations loaded with Fluo-4 AM (Fig. 2 A and B; Movie S1). A pulse of MC elicited a significant larger relative fluorescence response in mushroom-body regions of MC-trained females than in naive controls. Significantly enhanced responses were also observed in unexposed F1 and F2 descendants of trained females (Fig. 2C). These data associate the inherited behavioral bias with altered central processing of the same odor cue.

**Figure 2.**
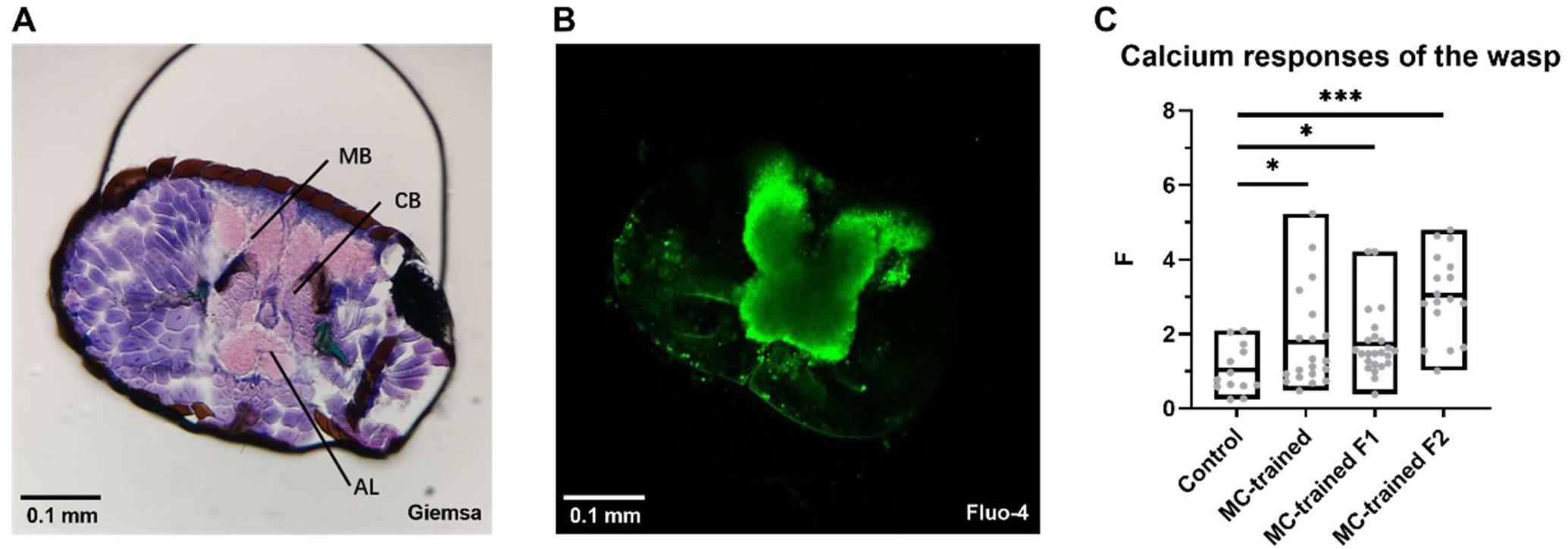
Anatomy and odor-evoked calcium activity in the wasp brain. (A) Transverse head section showing the mushroom body (MB), central brain (CB), and antennal lobe (AL). (B) Representative Fluo-4 signal in a semi-intact brain preparation. (C) Relative calcium responses in mushroom-body regions after MC stimulation in naive females, MC-trained females, and unexposed F1 and F2 descendants. Boxes show the observed range with the mean indicated. *P < 0.05;* P < 0.001.

### Reduced CCAPR expression is shared by trained females and descendants and is sufficient to shift odor choice

We compared head transcriptomes from naive females, females exposed to infected-host volatiles, females exposed to MC, and unexposed F1 descendants of MC-exposed females. Across the three experience-associated groups, 101 genes were differentially expressed relative to controls (Fig. S2). Enrichment analysis highlighted phosphatidylinositol signaling, including shared up-regulation of CDP-diacylglycerol-inositol 3-phosphatidyltransferase, diacylglycerol kinase eta, and the phosphatidylinositol 3-kinase regulatory subunit, consistent with altered neuronal signaling (Fig. S3).

Among neuropeptide-related genes, CCAPR was consistently down-regulated in females trained with infected-host volatiles or MC and in unexposed F1 descendants (Fig. 3A). Independent qPCR confirmed the reduction (Fig. 3B). Injection of CCAPR double-stranded RNA reduced CCAPR transcript abundance by 71.8% relative to the dsEGFP control and shifted odor choice away from MC and infected-host odor (Fig. 4 A-C). Thus, lowering CCAPR expression phenocopied the trained state and identifies CCAP signaling as a causal contributor to odor valuation.

**Figure 3.**
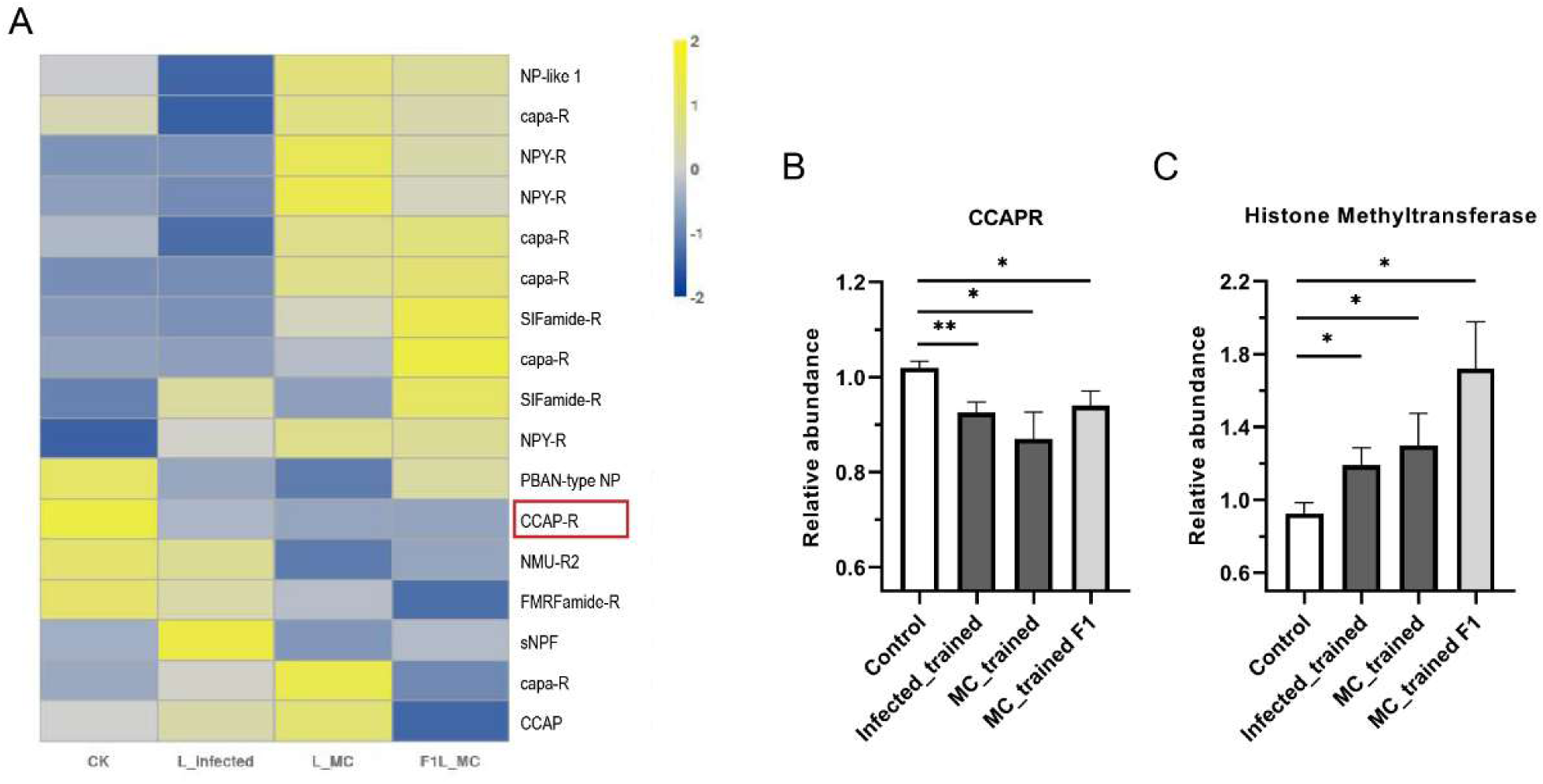
Transcriptomic signatures associated with trained and inherited odor avoidance. (A) Heat map of neuropeptide and neuropeptide-receptor transcripts. (B) Relative CCAPR transcript abundance. (C) Relative abundance of the histone methyltransferase candidate. The red box marks genes showing a shared expression trend across experience-associated groups. Data are means ± SE. *P < 0.05; P < 0.01;* P < 0.001.

**Figure 4.**
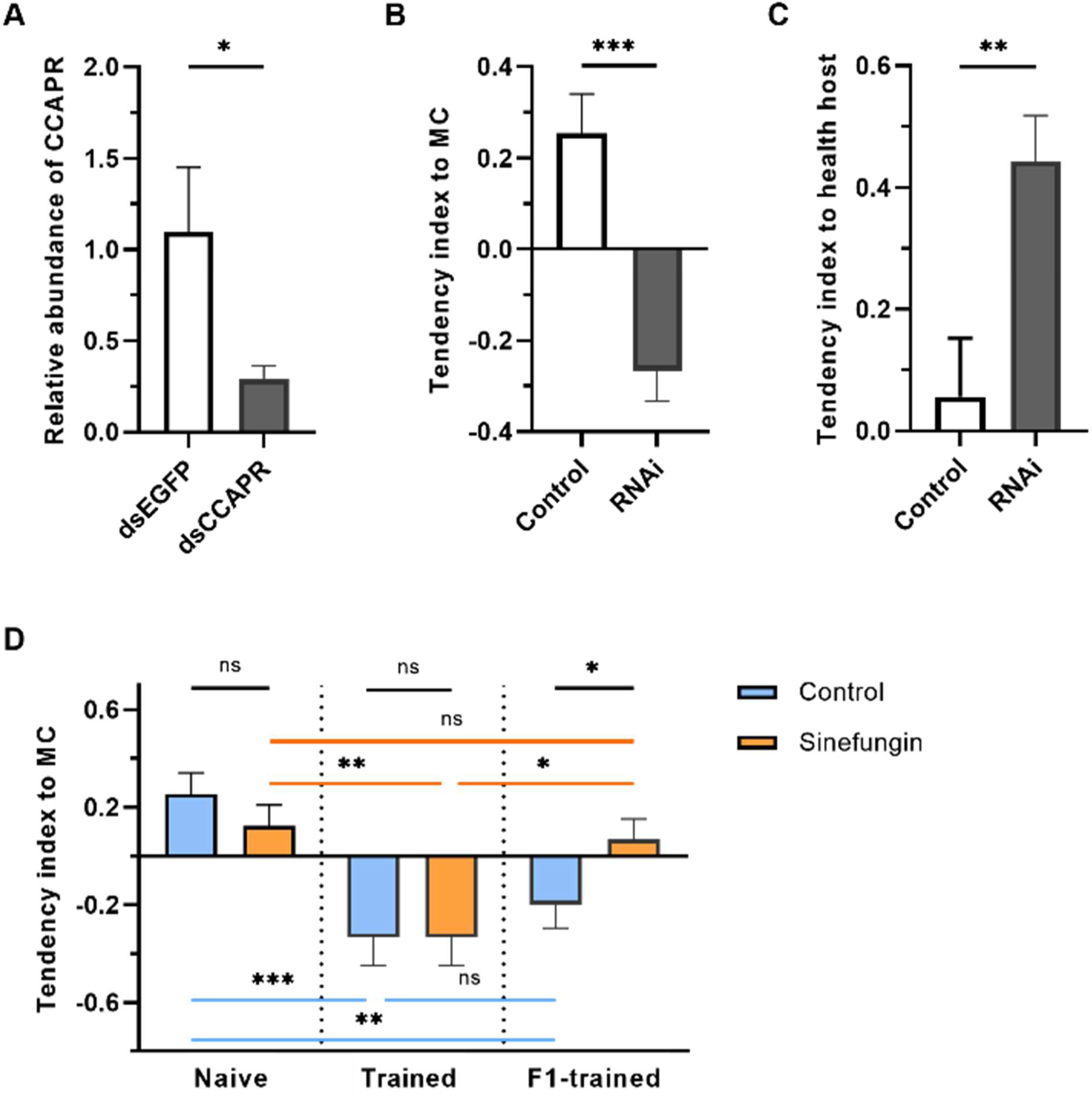
Effects of CCAPR RNAi and sinefungin on odor preference. (A) CCAPR knockdown efficiency. (B) Preference for MC versus solvent after dsCCAPR treatment. (C) Preference for healthy versus infected hosts after dsCCAPR treatment. (D) MC preference of naive and trained females treated with PBS or sinefungin and of their unexposed F1 offspring. Data are means ± SE. ns, not significant; *P < 0.05; P < 0.01;* P < 0.001.

### A sinefungin-sensitive methylation process is required for transmission but not maternal learning

A transcript annotated as a histone methyltransferase was up-regulated in trained females and in F1 descendants (Fig. 3C), prompting us to test whether methylation was required for the phenotype. Females were injected with sinefungin before MC exposure. Sinefungin did not alter baseline MC preference and did not prevent trained females from developing MC avoidance. In contrast, F1 offspring of sinefungin-treated, MC-exposed females no longer expressed the inherited avoidance bias (Fig. 4D). Because sinefungin inhibits multiple S-adenosylmethionine-dependent methyltransferases, these data support a methylation-sensitive transmission process but do not by themselves identify a specific histone mark or enzyme.

**Figure 5.**
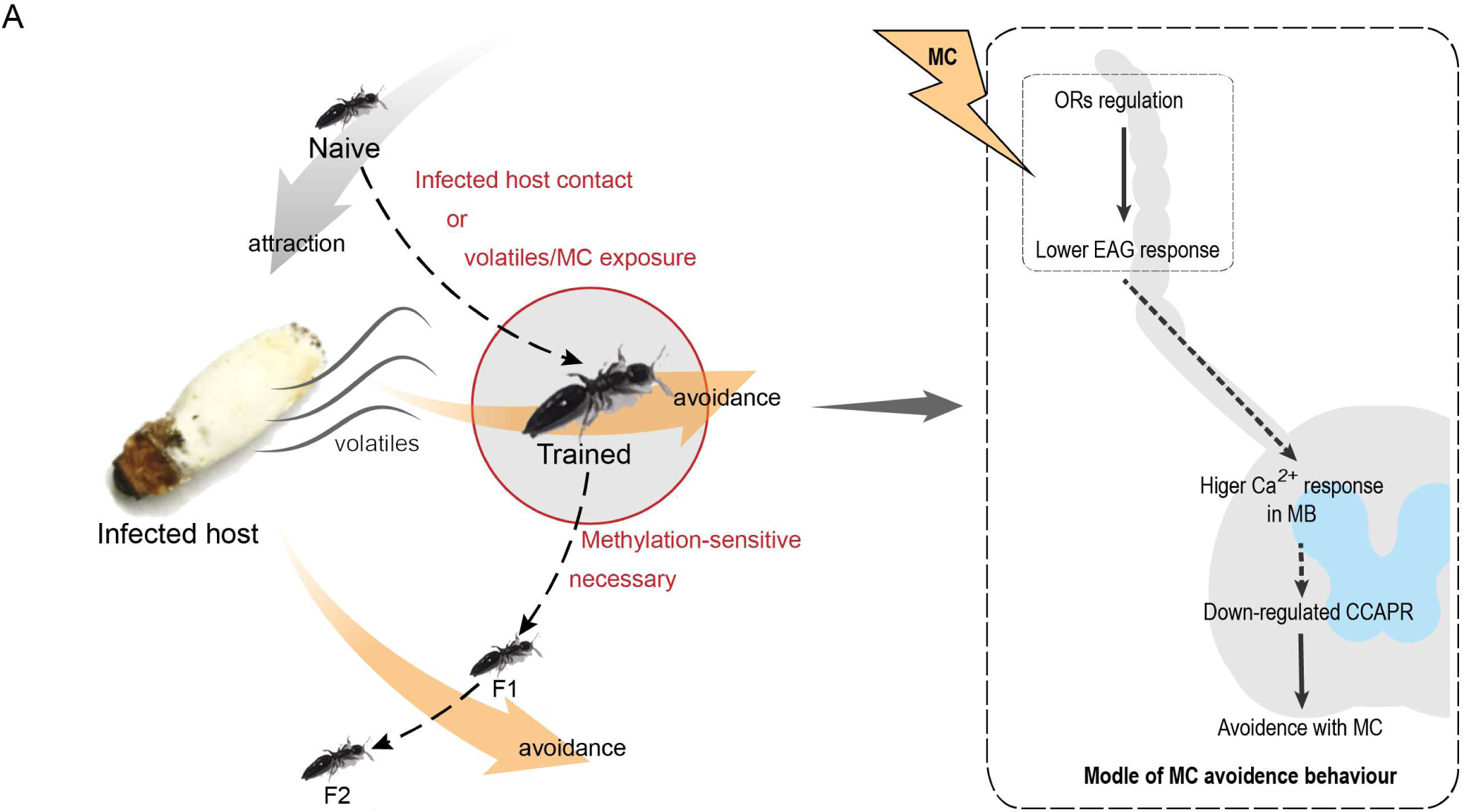
Working model for experience-dependent and inherited avoidance of an infection-associated odor. Prolonged exposure to infected-host volatiles or MC shifts behavior from attraction or indifference toward avoidance. The trained state is associated with reduced peripheral responses, enhanced mushroom-body activity, and reduced CCAPR expression. The behavioral bias persists in F1 and F2 descendants, and maternal sinefungin treatment blocks transmission to F1, implicating a methylation-sensitive process.

## Discussion

This study shows that prolonged experience with a pathogen-associated volatile changes the value assigned to that odor in a parasitoid wasp and that the resulting behavioral bias is expressed in two generations of unexposed descendants. The phenotype is accompanied by stronger odor-evoked activity in mushroom-body regions, convergent down-regulation of CCAPR, and loss of transmission after treatment of mothers with a broad methyltransferase inhibitor. Together, the results connect ecological sensory experience to central olfactory plasticity, neuropeptidergic state, and cross-generational behavioral change.

Prolonged odor exposure changes cue value, but the learning category should be defined cautiously. The noncontact experiment rules out the need for host handling, feeding, oviposition, or another obvious reinforcement and therefore distinguishes the effect from conventional host-associated conditioning. The most defensible description is non-associative olfactory plasticity. Classifying the response specifically as sensitization is less certain. Sensitization usually predicts enhanced responsiveness to the trained stimulus or related stimuli, whereas habituation predicts a reduced response (Anton et al., 2011; Minoli et al., 2012). Here, prolonged exposure increased behavioral avoidance, reduced antennal EAG amplitude, and increased mushroom-body calcium activity. Moreover, brief exposure increased attraction while prolonged exposure produced avoidance. This reversal suggests that at least two processes may operate over different timescales, potentially reflecting an initial resource-associated response followed by central reassignment of negative value. Similar duration-dependent or level-specific plasticity has been reported after pathogen odor exposure and chronic odor stimulation in other systems (Gugel et al., 2023; Pereira et al., 2020). Tests of generalization to unrelated odors, dishabituation, retention, and stimulus-response functions would be needed to distinguish sensitization, habituation, and other forms of non-associative plasticity.

Opposite peripheral and central changes suggest reweighting across the olfactory hierarchy. The reduction in EAG response after MC exposure indicates peripheral adaptation or altered receptor-neuron output, consistent with experience-dependent regulation of insect olfactory receptors (Claudianos et al., 2014). Yet MC evoked stronger calcium activity in the mushroom-body region of trained wasps and their descendants. In insect brains, mushroom-body circuits transform sparse odor representations into learned value and action bias, and their output can drive approach or avoidance (Aso et al., 2014; Peng and Chittka, 2017; Prisco et al., 2021; Séjourné et al., 2011). The combined EAG and imaging results therefore support central amplification or reweighting of an odor whose peripheral input is reduced. However, the present bulk calcium measurements do not resolve Kenyon cells, mushroom-body output neurons, or modulatory inputs, and stronger fluorescence should not be interpreted as a memory trace by itself. Cell-type-resolved imaging and simultaneous comparison of antennal-lobe and mushroom-body activity would establish where the inherited change first emerges.

CCAPR links neuropeptidergic state to odor-guided choice. CCAP is a conserved arthropod neuropeptide that regulates ecdysis, feeding, and foraging (Žitňan and Daubnerová, 2016). In flies, cockroaches, and beetles, CCAP interacts with NPF or sNPF pathways to alter foraging state and food responsiveness (Mikani et al., 2015; Shen et al., 2021; Williams, 2016). The shared reduction of CCAPR in trained females and descendants, together with the RNAi phenotype, extends this role to odor-guided host evaluation. The result does not yet establish that CCAPR stores odor-specific information. A parsimonious alternative is that reduced CCAP signaling changes the internal state through which the MC representation is evaluated. Rescue of CCAPR expression, localization of the relevant neurons, and tests with unrelated appetitive and aversive odors will be important for separating odor-specific memory from a broader change in motivation or response threshold.

The maternal effect contains a transgenerational component, but the transmitted entity remains unknown. The F1 response is an intergenerational maternal effect, whereas persistence to F2 after a single F0 exposure supports transgenerational inheritance under the breeding scheme used here (Sengupta et al., 2023). The finding parallels inherited odor biases in *Drosophila* and *Bicyclus* butterflies(Bozler et al., 2019; Dion et al., 2020; Gowri et al., 2019; Williams, 2016). Several mechanisms remain plausible: germline chromatin marks, small RNAs, altered maternal provisioning, neuroendocrine signals, microbial transmission, or direct transfer of odorant or odor-derived metabolites (Wang et al., 2017). The demonstration that hemolymph from odor-exposed butterfly larvae can transfer both an acquired preference and an offspring effect makes the last two possibilities especially relevant (Gowri and Monteiro, 2024). Reciprocal crosses, family-resolved breeding, analysis of ovaries and eggs, and chemical measurements of MC or its metabolites would distinguish germline information from maternal carryover.

The sinefungin experiment narrows the mechanism but should not be described as proof of histone methylation. Sinefungin is a S-adenosylmethionine analog that inhibits RNA, DNA, and protein methyltransferases as well as histone-directed enzymes (Yebra et al., 1991). Its selective effect on F1 transmission, together with increased expression of a histone methyltransferase candidate, supports a methylation-sensitive step between maternal experience and offspring phenotype. It does not identify the substrate, the tissue, or the developmental window involved. Direct measurement of candidate histone marks in ovaries or embryos, enzyme-specific knockdown, and rescue experiments are required. Studies in honeybees and nematodes demonstrate that DNA and histone methylation can modulate learned odor processing and inherited physiology, providing tractable hypotheses for follow-up (Biergans et al., 2012; Wang et al., 2017).

The infected-host odor is informative only when parental and offspring environments are correlated. Since the efficacy of biocontrol agents is strongly limited by environmental conditions and their dispersal, combining different species of agents is explored as solutions to counteract ecological constrains (Roy and Pell, 2000; Pobozniak and Olczyk, 2025). In such scenario, inherited avoidance could reduce costly attacks on unsuitable infected hosts and facilitate successful reproduction of the parasitoid population; when environments change, the same bias could exclude usable resources.

From an applied perspective, odor pre-exposure might eventually be used to alter host selection in mass-reared parasitoids deployed with entomopathogenic fungi. Since several studies explored the combined application of entomopathogens and *S. guani* against beetles (Li et al., 2017; Liu et al., 2007; Xu et al., 2014), the preference to healthy hosts would benefit to the overall biocontrol efficiency as well as long-term pest management success. Compared with associate learning that train wasps to associate odor through feeding or oviposition experiences, non-associative learning possess the advantage of convenient and feasible operation. Previous studies have largely concentrated on the conditioning of plant volatiles to improve host location and manipulating host species bias (Giunti et al., 2015), while this study provided a new perspective on the utilization of learning to train parasitoids to avoid entomopathogens infected hosts. That possibility should remain a secondary implication until experiments show improved parasitism, reproduction, and population persistence under realistic field conditions and exclude trade-offs in responses to other host cues.

## Conclusion

A purely olfactory experience with a pathogen-associated cue can durably alter odor-guided behavior in *S. guani* and bias the behavior of unexposed descendants. The phenotype combines attenuated peripheral responses, enhanced mushroom-body activity, reduced CCAPR expression, and methylation-sensitive maternal transmission. Framed as a problem of inherited sensory plasticity rather than pest-control optimization, the study provides a broadly relevant example of how environmental information may be converted into behavioral state across generations. The data support non-associative olfactory plasticity and a transgenerational component, while the precise learning process and molecular carrier remain important targets for future work.

## Materials and Methods

### Insects and preparation of infected hosts

A laboratory colony of *S. guani* was maintained on final-instar larvae of *Monochamus alternatus* for more than 90 generations. Host larvae were collected in Zhejiang Province, China, and stored individually in perforated 15-mL tubes containing moist sawdust at 4 °C. For colony maintenance, a mated female wasp was introduced to one host larva in a glass tube and held at 26 ± 2 °C and 70% relative humidity in darkness until adult offspring emerged. Newly emerged adults were stored at 4 °C for no longer than 2 months and reactivated at 26 °C for 24 h before experiments. To produce infected hosts, larvae weighing 250-350 mg were surface-sterilized with 75% ethanol and immersed for 8-10 s in a *B. bassiana* spore suspension prepared from BotaniGard 22WP (1 × 10^9 conidia mL^-1 in 5% Tween 20). Treated larvae were held individually at 26 ± 2 °C and 70% relative humidity. Hosts covered by dense, moist mycelium 6-7 d after inoculation were used in training and choice assays.

### Odor-choice assays

Odor preference was measured in a Y-tube olfactometer. Females chose between a healthy and an infected host or between hexane and MC (10 μg mL^-1 in hexane). For chemical-choice assays, 5 μL of each solution was applied to a circular filter paper (0.8-cm diameter). Each female was tested once, and the olfactometer was ventilated between trials. Choice outcomes were summarized in groups of five females as a healthy-host tendency index, (number choosing healthy host - number choosing infected host)/5, or an MC tendency index, (number choosing MC - number choosing hexane)/5.

For the free-choice assay, 60 females were released into a 10-cm Petri dish containing two foil-wrapped glass tubes with perforated openings. One tube contained an infected host and the other was empty. After 24 h, a choice was recorded when a female entered one of the tubes.

### Odor exposure and production of descendant generations

For contact exposure, groups of 10 females were confined with one infected host in a glass tube for 12 h at 26 ± 2 °C and 70% relative humidity in darkness. For noncontact exposure, individual females were confined in ventilated glass tubes that prevented physical contact with the host. Twenty female-containing tubes were oriented toward four infected hosts inside a sealed 15-cm Petri dish and exposed for 12 h under the same environmental conditions.

For MC exposure, the arrangement was identical except that cotton balls loaded with 20 μL MC solution (10 μg mL^-1 in hexane) served as odor sources. After exposure, females were held individually at 25 ± 2 °C, and odor preference was tested within 24 h. In the duration experiment, females were exposed to infected-host volatiles for 0.5, 1, 9, or 12 h.

For generation experiments, exposed females were not behaviorally tested. Twelve hours after exposure, females were introduced to hosts and allowed to reproduce under colony conditions. Their offspring were designated F1. Unexposed F1 females were allowed to reproduce to generate F2. F1 and F2 females were reactivated under the same conditions as the parental generation before odor-choice assays.

### Histology and calcium imaging

For anatomical sections, heads were incubated overnight in 20 g L^-1 sucrose, transferred for 2 h to a 1:1 mixture of sucrose solution and optimal cutting temperature compound, incubated for 4 h in pure compound, and then transferred to fresh compound for 6 h. Heads were sectioned at 10 μm at −22 °C and stained with 5% Giemsa for 30 min.

Odor-evoked activity was measured in semi-intact head preparations loaded with Fluo-4 AM. MC-trained females were imaged immediately after exposure; naive females and descendants were reactivated at 26 °C for 36 h before imaging. A female was restrained in a 10-μL pipette tip with the antennae exposed. The posterior third of the head was removed, and the preparation was maintained in insect Ringer solution. Preparations were incubated for 1 h in darkness in 200 μL loading solution containing 3 μL 0.2 mM Fluo-4 AM, 1 μL 20% Pluronic F-127 in DMSO, and 196 μL Ringer solution, then returned to Ringer solution and imaged within 12 h.

Fluorescence was recorded with a Nikon A1R MP two-photon microscope using 920-nm excitation at one frame per second. For odor delivery, 5 μL MC was applied to filter paper in a 5-mL syringe and allowed to equilibrate for at least 3 min. A 1-mL pulse of headspace was delivered from approximately 10 cm at 15 s in a 30-s recording. The imaging area was ventilated between trials. Regions of interest in mushroom-body terminal regions were traced in ImageJ. The peak fluorescence after stimulation was Ft, baseline fluorescence 1 s before the peak was F0, and the relative response was calculated as ΔF/F0 = (Ft - F0)/F0.

### RNA sequencing and quantitative PCR

Four groups were analyzed: naive controls (CK), females exposed to infected-host volatiles (L_infected), females exposed to MC (L_MC), and unexposed F1 descendants of MC-exposed females (F1L_MC). Each biological replicate contained 40 pooled heads, with three replicates per group. CK and F1L_MC females were collected after 36 h of reactivation at 26 °C; exposed females were collected immediately after training. Samples were frozen in liquid nitrogen, and heads were separated on dry ice and stored at −80 °C.

Total RNA was extracted with TRIzol. Poly(A)+ RNA was enriched with oligo(dT) magnetic beads, fragmented, converted to double-stranded cDNA, end-repaired, adapter-ligated, amplified, and purified using an Illumina-compatible library protocol. Libraries were quality checked on an Agilent 2100 Bioanalyzer and sequenced as 150-bp paired-end reads on an Illumina NovaSeq platform.

Reads were aligned with HISAT2 v2.0.5, and gene counts were generated with featureCounts v1.5.0. Expression was summarized as FPKM, and differential expression was tested with DESeq2 v1.20.0. Genes with Benjamini-Hochberg adjusted P ≤ 0.05 were considered differentially expressed. KEGG enrichment was analyzed with clusterProfiler.

For qPCR, 1 μg total RNA was reverse-transcribed with the TIANScript kit. Reactions were run with Taq Pro Universal SYBR qPCR Master Mix on an ABI QuantStudio 12K system. Primer sequences and reference genes are listed in Supplementary Table S1.

### RNA interference

Double-stranded RNA targeting CCAPR and EGFP was synthesized with the T7 RiboMAX Express RNAi System using primers in Supplementary Table S1. After 24 h of reactivation, females were chilled on ice and injected between the second and third abdominal segments with 80 ng dsCCAPR or dsEGFP in 20 nL using a Micro4 microsyringe pump. Females were held individually for 48 h before qPCR and odor-choice assays.

### Methyltransferase-inhibitor treatment

Females were injected with 20 nL of 0.05 M sinefungin in PBS (1 nmol per female); control females received 20 nL PBS. After 6 h at 26 ± 2 °C and 70% relative humidity in darkness, females were exposed to MC. One subset was tested for MC preference, and a second subset was allowed to reproduce. MC preference was then measured in their unexposed F1 offspring.

### Statistical analysis

Analyses were performed in GraphPad Prism 8. Normality was assessed with the Shapiro-Wilk test. Parametric two-group comparisons used Student’s t test, with Welch’s correction when variances were unequal. Nonparametric comparisons used the Kruskal-Wallis test. Variance homogeneity was assessed with F tests or Bartlett’s tests as appropriate. Statistical significance was set at P < 0.05. Exact sample sizes, test statistics, and source data should be reported for every panel in the final submission.

## Acknowledgments

This work was funded by grants from the National Natural Science Foundation of China (32088102, 32061123002), Hebei Natural Science Foundation (C2022201042), and High-level Talent Research Funding Project of Hebei University (070006-70280052401).

## Supplementary Figure Legends and Figures

**Movie S1. Example movie of calcium-imaging acquisition and MC stimulation.**

**Table S1.**
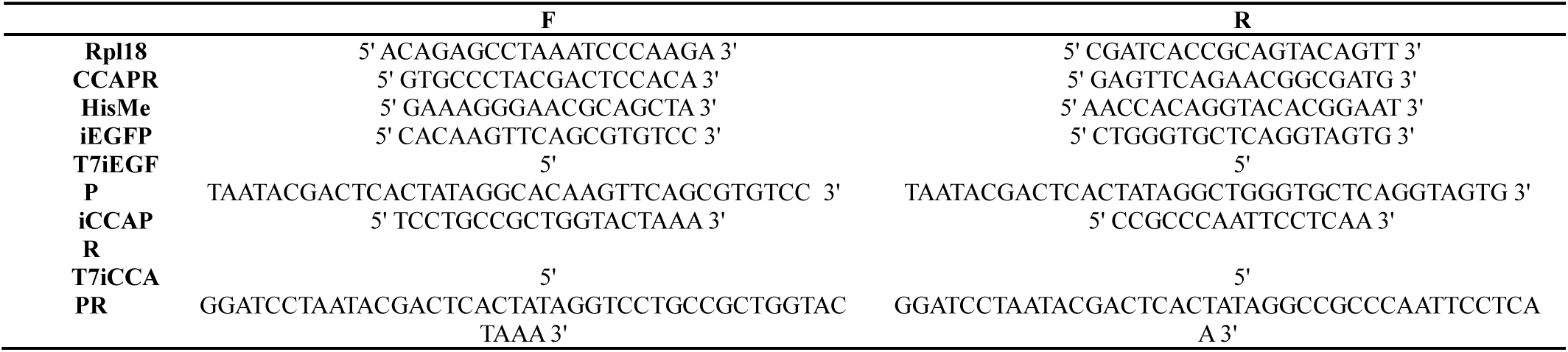
Primers.

**Figure S2.**
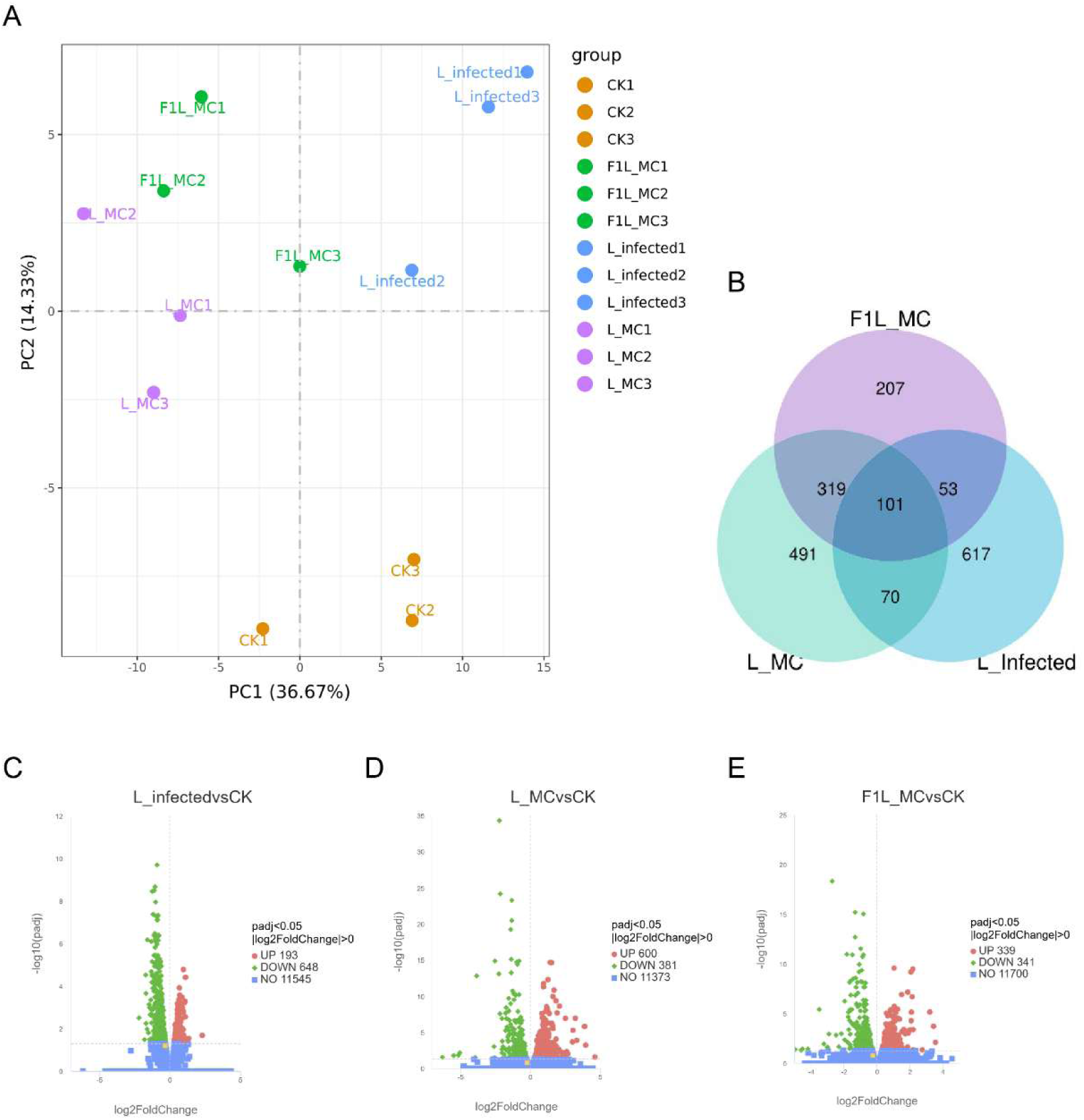
Transcriptome overview. (A) Principal-component analysis. (B) Venn diagram of differentially expressed genes across the three experience-associated comparisons. (C-E) Volcano plots for L_infected versus CK, L_MC versus CK, and F1L_MC versus CK. Orange points indicate CCAPR.

**Figure S3.**
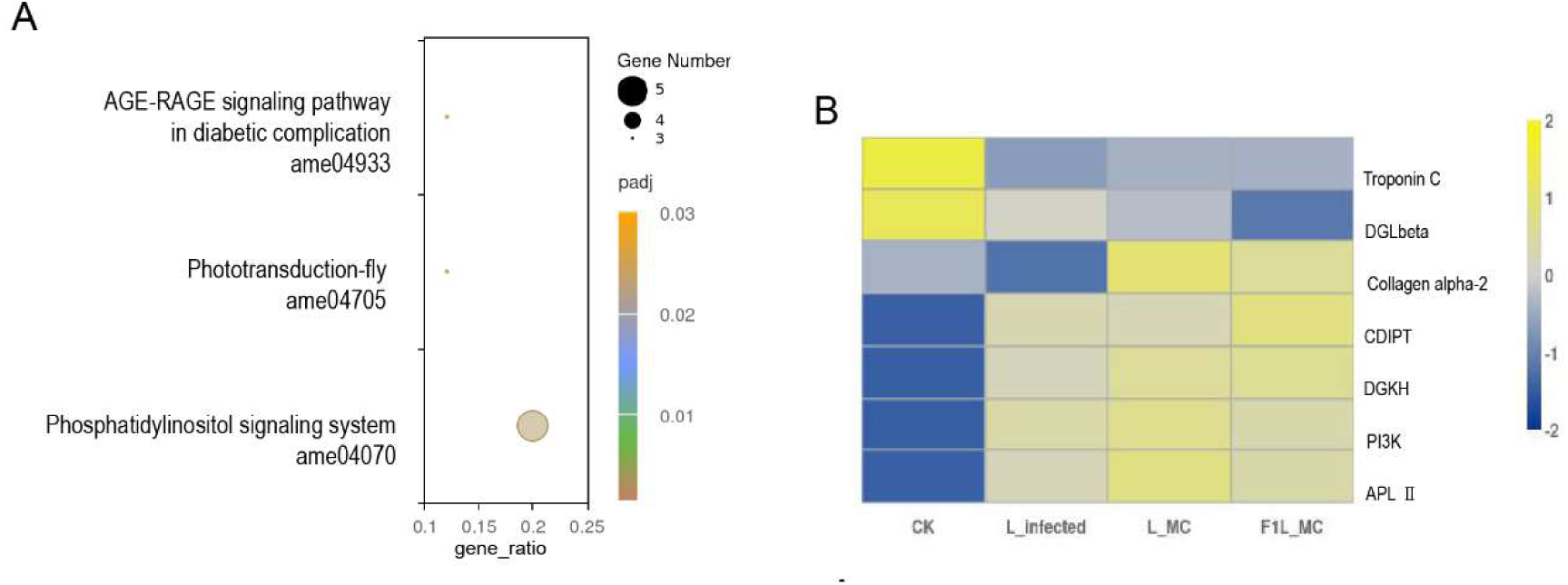
KEGG enrichment of shared differentially expressed genes. (A) Enriched KEGG pathways among the 101 shared differentially expressed genes. (B) Heat map of genes contributing to enriched pathways.

**Figure S4.**
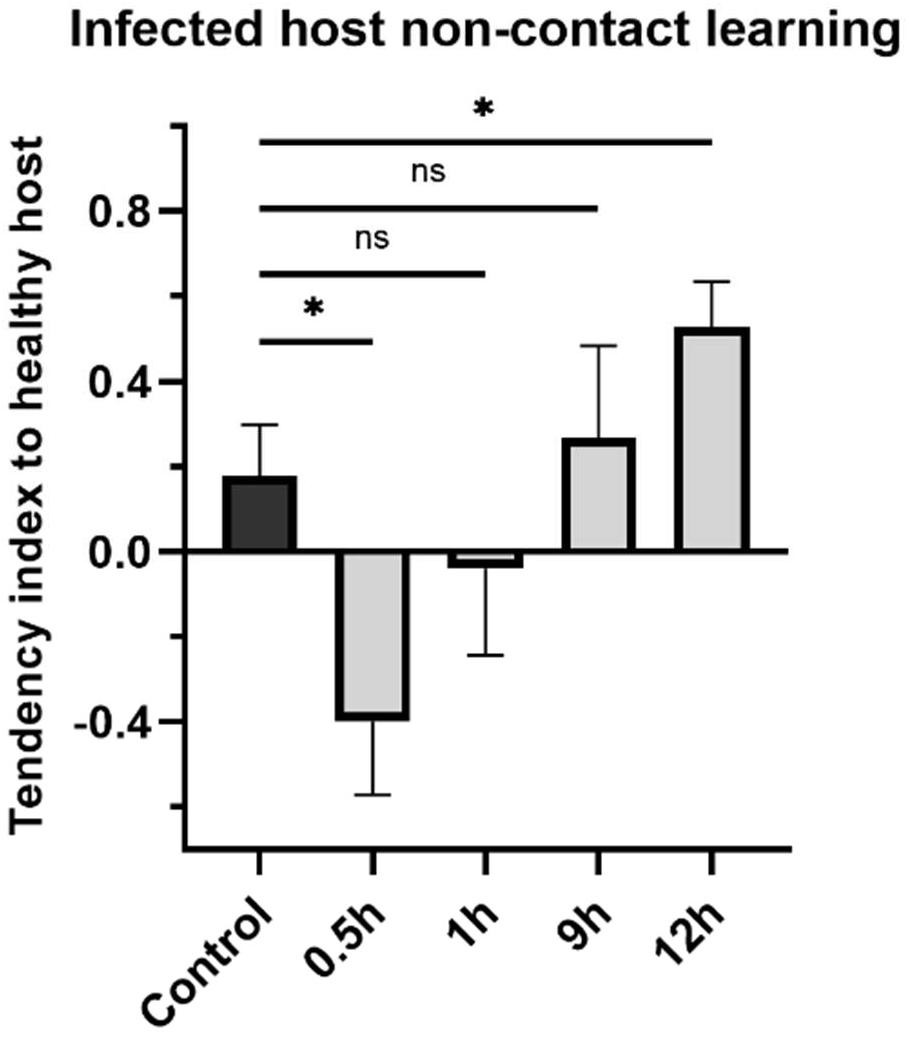
Duration-dependent effect of infected-host odor exposure. Preference for healthy versus infected hosts after 0.5, 1, 9, or 12 h of noncontact exposure. Data are means ± SE. ns, not significant; *P < 0.05.

**Figure S5.**
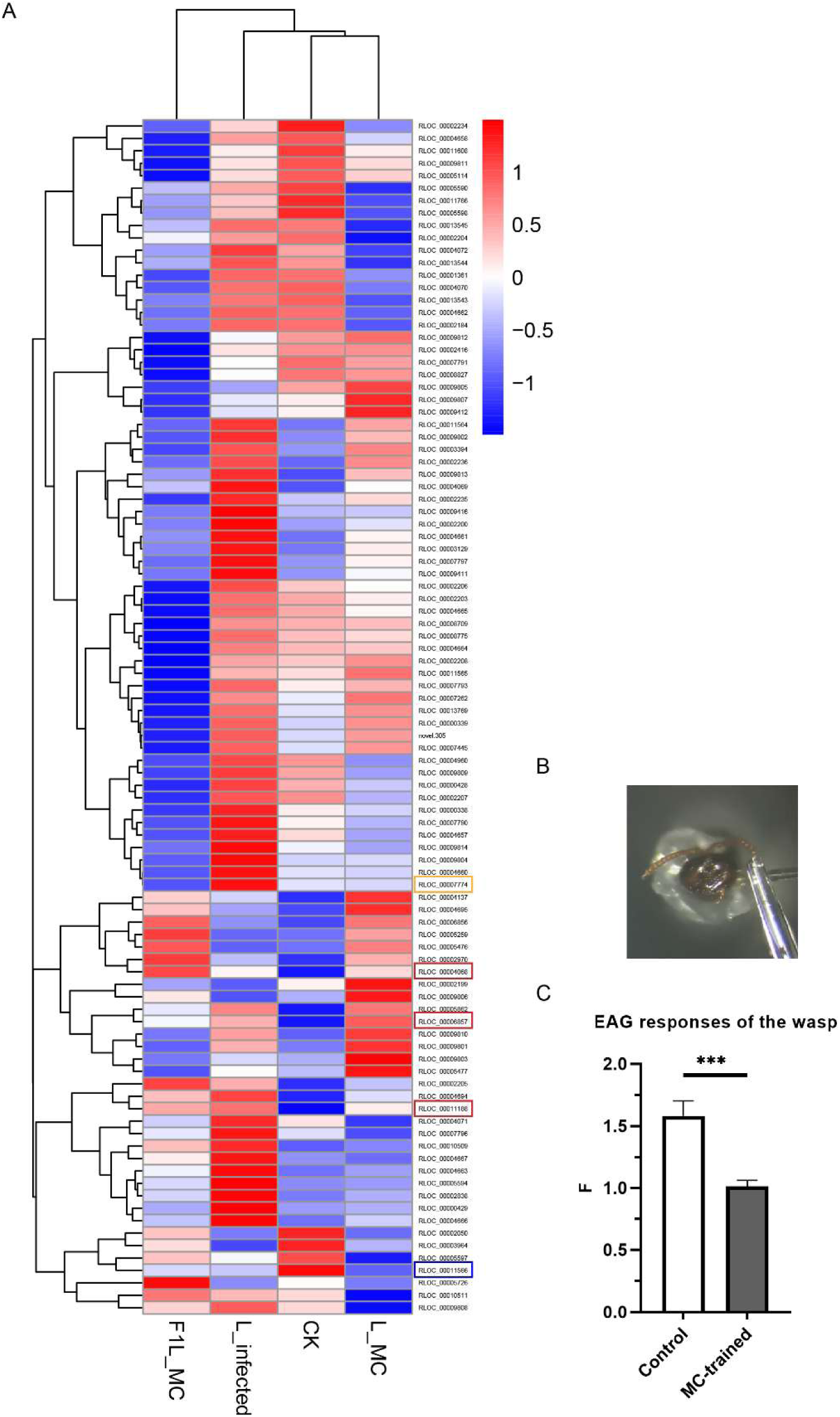
Olfactory-receptor expression and antennal responses to MC. (A) Heat map of odorant-receptor transcripts. (B) Representative electroantennogram preparation. (C) EAG responses to MC in naive and MC-trained females (n = 9 per group). Colored boxes mark receptor genes with shared expression trends; the orange box marks SguaOR48. Data are means ± SE. *P < 0.001.

Data S3. Raw data of behaviour and molecular experiments

## Data Availability

All data generated and analyzed in this study are provided in the Supplementary Information and associated source-data files. Sequence data are deposited in a public repository (https://ngdc.cncb.ac.cn/gsa/s/4e439NUC).

